# B-Myb association with DNA is mediated by its negative regulatory domain and Cdk phosphorylation

**DOI:** 10.1101/2022.06.13.495963

**Authors:** Tilini U. Wijeratne, Keelan Z. Guiley, Hsiau-Wei Lee, Gerd A. Müller, Seth M. Rubin

## Abstract

B-Myb is a highly conserved member of the vertebrate Myb family of transcription factors that plays a critical role in cell-cycle progression and proliferation. Myb proteins activate Myb-dependent promoters by interacting specifically with Myb binding site (MBS) sequences using their DNA binding domain (DBD). Transactivation of MBS promoters by B-Myb is repressed by its negative regulatory domain (NRD), and phosphorylation of the NRD by Cdk2-CyclinA relieves the repression to activate B-Myb dependent promoters. The structural mechanisms underlying autoinhibition and activation have been poorly characterized. We determined that a region in the B-Myb NRD (residues 510-600) directly associates with the DBD and inhibits DBD binding to the MBS DNA sequence. We demonstrate that phosphorylation of the NRD at T515, T518, and T520 is sufficient to disrupt the interaction between the NRD and the DBD, which results in increased affinity for MBS DNA and increased B-Myb-dependent promoter activation. Our biochemical characterization of B-Myb autoregulation and the activating effects of phosphorylation provides insight into how B-Myb functions as a site-specific transcription factor.

## Introduction

The Myb family of transcription factors (TFs) are present in a range of species from slime mold to higher eukaryotes and have high conservation in their DNA binding domain (DBD) (1-4). TFs with evolutionary conserved DBDs recognize a common DNA sequence; however, they often diverge in their distinct functions through different intra- and inter-molecular interactions (5). Vertebrate Myb family members A-Myb, B-Myb, and c-Myb share more than 70% amino sequence homology in their DBDs, which recognize the Myb binding site (MBS) DNA sequence (C/TAACNG) (4,6-8). All the Myb family members regulate transcription of genes important for cell differentiation and proliferation, but they differ in their tissue-specific expression. A-Myb (encoded by *MYBL1)* is mainly expressed in cells of the developing central nervous system, sperm cells, and breast tissue, while c-Myb (*MYB*) is expressed specifically in immature hematopoietic stem cells (9,10). In contrast, B-Myb (*MYBL2)*, which is the most ancient of the paralogs, is ubiquitously expressed in all proliferating cells (11). The prominent role of B-Myb in both differentiating and proliferating cells is reflected by its deregulation in several cancers. Over-expression of the *MYBL2* gene is considered a biomarker for poor prognosis in osteosarcoma, breast cancer, esophageal cancer, and multiple myeloma (12-15). Therefore, in recent years B-Myb has become an attractive target to understand mechanistic details of oncogenic TFs for cancer therapeutics.

The B-Myb domain architecture is similar to A-Myb and c-Myb. B-Myb contains a DBD, a negative regulatory domain (NRD) toward the C-terminus, and a transactivation domain (TAD) (Fig. 1A). In all three Myb proteins, C-terminal protein truncations that remove the NRD trigger activation of Myb-dependent reporter promoters (16-18). Recurrent chromosomal translocations involving the genes *MYB* and *MYBL1* produce NRD truncated versions of the respective proteins c-Myb and A-Myb that are sufficient to induce leukemias in mice (19,20). These truncated proteins more resemble the viral oncoprotein v-Myb, which shares the same DBD as all the Myb proteins and has a TAD but lacks a potent C-terminal NRD (21). In contrast, C-terminal truncations of B-Myb are not reported to have oncogenic properties. However, the B-Myb C-terminus contains the MuvB binding domain (MBD), which binds the MuvB complex to assemble Myb-MuvB (MMB) (22-24). The MMB complex activates cell-cycle dependent genes that contain a CHR sequence (cell-cycle genes homology region) in their promoter in a manner that is both B-Myb and MuvB dependent (24). Thus, B-Myb functions as a site-specific TF that activates MBS genes and as a co-activator of CHR genes when present in the MMB complex. This latter function is unique among Myb family members.

**Figure 1.**
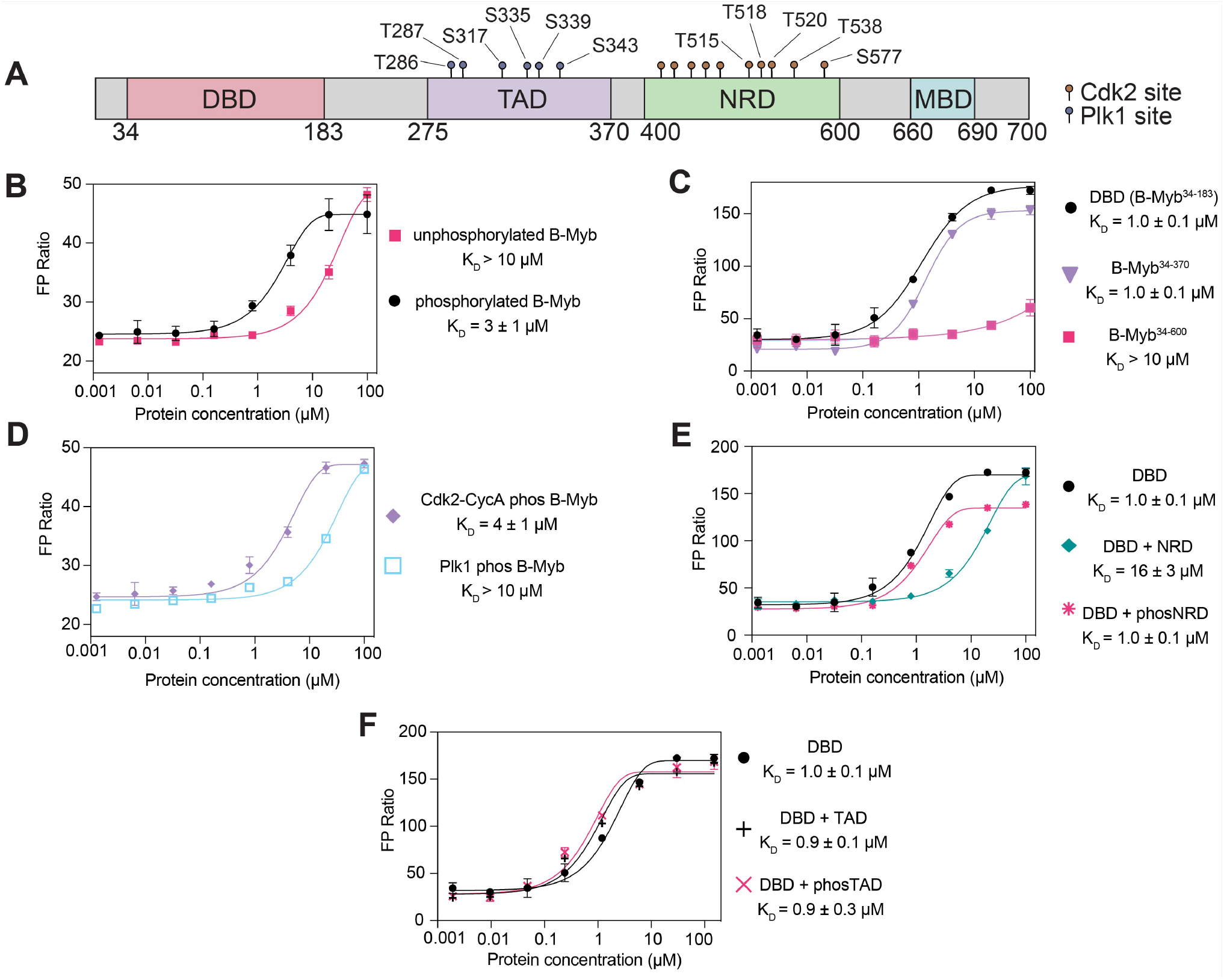
Phosphorylation by Cdk2-CycA enhances B-Myb binding to MBS DNA. (A) A schematic presenting the boundaries of DBD the (DNA binding domain), TAD (transactivation domain), NRD (negative regulatory domain), and MBD (MuvB binding domain). Previously identified Cdk2 and Plk1 sites are indicated. (B) Fluorescence polarization (FP) assay of B-Myb binding to a TAMRA-labeled MBS probe. The measurements compare unphosphorylated B-Myb to B-Myb that has been sequentially phosphorylated by Cdk2-CyclinA and Plk1. (C) Same FP assay used to measure binding affinities of C-terminal truncations of B-Myb to the MBS probe. (D) Same FP assay used to measure probe affinity of B-Myb phosphorylated only by Cdk2-CyclinA or Plk1. (E) FP assay of DBD binding to MBS probe as in panel C but also performed by using DBD incubated with 30 μM NRD or phosNRD (F) As in panel E but using DBD incubated with 30 μM TAD or phosTAD.

The mechanisms by which the NRD affects the transactivation potential of B-Myb are also not yet fully understood. Several studies show that autoinhibition of B-Myb by the NRD is relieved when B-Myb is phosphorylated by the cell-cycle regulatory kinase Cdk2-CyclinA (Cdk2-CycA), which results in activation of MBS dependent promoters (17,25,26). The NRD of Myb proteins contains several highly conserved Cdk consensus sites (S/TP) (Fig. 1A and Fig. S1), including a TPTPFK motif (amino acids 519-524 in B-Myb). This region is a direct target of Cdk-mediated phosphorylation in cell-based studies and shows a positive correlation with activation (27,28). Cdk-dependent phosphorylation of B-Myb also primes for binding of Polo-like kinase 1 (Plk1), and subsequent phosphorylation by Plk1 in the TAD also promotes B-Myb activity (29). Although evidence suggests that NRD phosphorylation releases an inhibited state of B-Myb to significantly transactivate MBS-dependent promoter reporters, two studies did not find that Cdk2-CycA alters B-Myb interactions with DNA (30,31). However, evidence of enhanced DNA binding upon truncation of the NRD in A-Myb and c-Myb has been reported (18,27). A recent study showed that the B-Myb DBD undergoes an intramolecular interaction with NRD. Cdk-mediated phosphorylation at a specific site (S577) disrupted the interaction; however, it was not conclusive if the NRD-DBD interaction affected the ability of B-Myb to bind DNA and if phosphorylation regulated the NRD-DBD association (31). B-Myb phosphorylation is not only required for MBS-dependent transactivation, but it is also important for G2/M cell cycle-dependent gene activation. B-Myb is extensively phosphorylated by Cdk2-CycA during S phase of the cell cycle coinciding with its peak in expression (24,29). On the other hand, other studies conclude that extensive phosphorylation of B-Myb is important for its ubiquitination and proteasomal mediate degradation (32). Thus, despite these various studies, it remains unclear how phosphorylation of B-Myb overcomes negative regulation by the NRD to activate B-Myb. In part, disparate models have arisen because of the challenges of interpreting cell-based assays to draw conclusions about direct molecular interactions and the effects of phosphorylation on specific interactions.

Here, we present a study of B-Myb autoregulation that focuses on biophysical assays using purified proteins. We used bacterial recombinant expression, which provided us with a minimal system to control phosphorylation, and we quantified interactions by measuring dissociation constants. We found that the B-Myb NRD binds the DBD with a low micromolar affinity, and the interaction is sufficient to inhibit B-Myb binding to MBS DNA. We identified amino acids that are critical for NRD-DBD association and observed that Cdk2-CycA mediated phosphorylation of T515, T518, and T520 disrupts the NRD-DBD interaction to enhance binding to an MBS DNA sequence probe. We also show that specific mutations that disrupt the NRD-DBD interaction increase B-Myb-dependent activation of an MBS luciferase reporter. Our findings reveal a structural mechanism for B-Myb autoregulation of site-specific gene activation and how repression is relieved by Cdk phosphorylation.

## Results

### Phosphorylated B-Myb binds to DNA tighter than unphosphorylated B-Myb

In order to determine the effects of B-Myb phosphorylation on its association with DNA, we used fluorescence polarization (FP) assays to measure binding affinities using recombinant, purified proteins. Considering that both Cdk2-CycA and Plk1 sites are present throughout the NRD and TAD regions respectively (Fig. 1A and Fig. S1) (27,29,30), we sequentially phosphorylated purified, full-length B-Myb. We phosphorylated first with Cdk2-CycA and then with Plk1, and we verified phosphorylation with a mobility shift on a PhosTag gel (Fig. S2A-S2B). We then assayed DNA binding of phosphorylated and unphosphorylated protein to a fluorescently labeled DNA probe containing the Myb binding site (MBS) sequence. The resulting single-site protein-DNA binding curve for the phosphorylated B-Myb showed a dissociation constant K_D_ = 3 ± 1 µM (Fig. 1B). The unphosphorylated B-Myb showed weak (K_d_ > 10 µM), potentially nonspecific, binding to the probe.

### Phosphorylation of the NRD regulates DBD binding to DNA

We found that the affinity of phosphorylated B-Myb for the MBS probe is similar to the affinity of the DBD alone (B-Myb^34-183^, K_D_ = 1.0 ± 0.1 µM) and to the affinity of a construct in which the NRD is deleted (B-Myb^34-370^, K_D_ = 1.0 ± 0.1 µM) (Fig. 1C). In contrast, a C-terminal truncation of the MBD that leaves the NRD intact (B-Myb^34-600^) binds with similar weak affinity as unphosphorylated full-length B-Myb (Fig. 1C). The observation that binding of the construct containing the DBD-TAD-NRD domains is greatly reduced compared to both the DBD and DBD-TAD only constructs demonstrates that the NRD inhibits DBD binding to the MBS probe. These results and the fact that there are no Cdk sites in the DBD support a model in which the DBD-DNA interaction is inhibited by the NRD in the context of unphosphorylated B-Myb and that this inhibition is relieved upon Cdk2-CycA mediated phosphorylation of the NRD. To test this model, we phosphorylated B-Myb with Cdk2-CycA only and found a similar affinity as phosphorylating with both kinases. In contrast, phosphorylation with Plk1 only had no effect on the binding compared to unphosphorylated B-Myb (Fig. 1D and Fig. S2).

To further test the role of the NRD in inhibiting DBD-DNA binding and the role of NRD phosphorylation, we performed FP assays, titrating DBD into DNA in the presence of unphosphorylated and Cdk2-phosphorylated NRD^440-600^ (verified by mass spectrometry, Fig. S2C). As shown in Fig. 1E, when added *in trans*, 30 µM unphosphorylated NRD reduced DBD binding to the MBS probe (K_D_ = 16 ± 3 µM). In contrast, when 30 µM phosphorylated NRD was added *in trans*, there is little effect of the NRD on the affinity of the DBD for the MBS probe (K_D_ = 1.0 ± 0.1 µM). The addition of Plk1-phosphorylated TAD *in trans* also did not influence DBD binding to the MBS probe (Figs. 1F and S2D). These data further support the model that Cdk phosphorylation of NRD^440-600^ and not Plk1 phosphorylation of sites in the TAD increases the affinity of DNA binding through release of autoinhibition.

### Direct Association of the NRD with the DBD

We next probed the presence of intramolecular interactions within B-Myb that may drive the observed autoinhibition of DNA binding. We mixed separately purified domains and detected interdomain association by isothermal titration calorimetry (ITC). We observed no detectable binding of the TAD or MBD to an NRD construct that that contains amino acids 440-600 (NRD^440-600^, Figs. 2A and 2B). In contrast, we detected association of the DBD and NRD^440-600^ and measured an affinity of K_D_ = 4.5 ± 0.5 µM (Fig. 2C). To map a more minimal NRD, we used sequence conservation to divide the NRD into two halves (Fig. S1). We observed that the C-terminal half (amino acids 510-600, B-Myb^510-600^) binds the DBD with a similar affinity of K_D_ = 4.9 ± 0.2 µM (Fig. 2D), and we did not observe association of the N-terminal half (B-Myb^440-510^, Fig. 2E).

**Figure 2.**
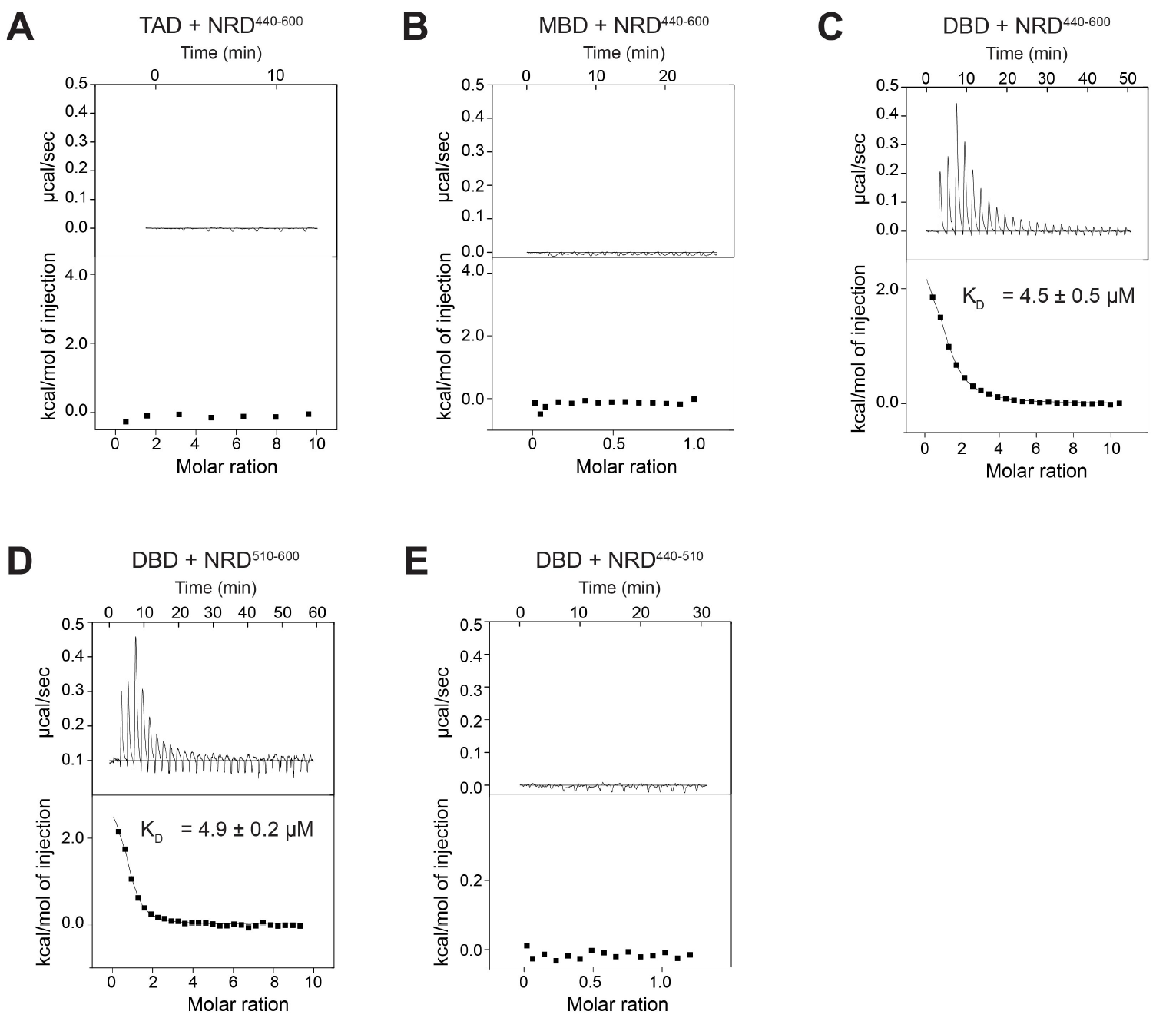
B-Myb NRD^510-600^ directly binds DBD. (A) ITC binding measurement between NRD^440-600^ and TAD. (B) ITC binding measurement between NRD^440-600^ and MBD. (C) Isothermal titration calorimetry (ITC) binding measurement between NRD^440-600^ and DBD. (D) ITC binding measurement between NRD^510-600^ and DBD. (E) ITC binding measurement of NRD^440-510^ and DBD.

### NMR spectroscopy maps amino acids involved in the B-Myb DBD-NRD interaction

We next used nuclear magnetic resonance (NMR) to further probe the NRD^510-600^-DBD interaction. The minimal chemical shift dispersion of the ^1^H-^15^N HSQC spectrum of ^15^N-labeled B-Myb^510-600^ suggests that the fragment is structurally disordered (Fig. 3A). We therefore generated a ^13^C-^15^N double-labeled sample and proceeded with ^13^C-^15^N CON spectroscopy, which is well suited for studying interactions of intrinsically disordered proteins (33). To observe the NRD-DBD association, we added isotopically unlabeled DBD to ^13^C-^15^N labeled NRD and monitored the chemical shift perturbations in a two-dimensional CON spectrum. A number of cross-peaks showed changes in both intensity and position, which is consistent with the binding we observed by ITC (Fig. 3B). Most of the perturbations appear as loss of intensity, which reflects peak broadening from either intermediate exchange or from the NRD^510-600^ forming a larger molecular weight complex when bound by the unlabeled DBD.

**Figure 3.**
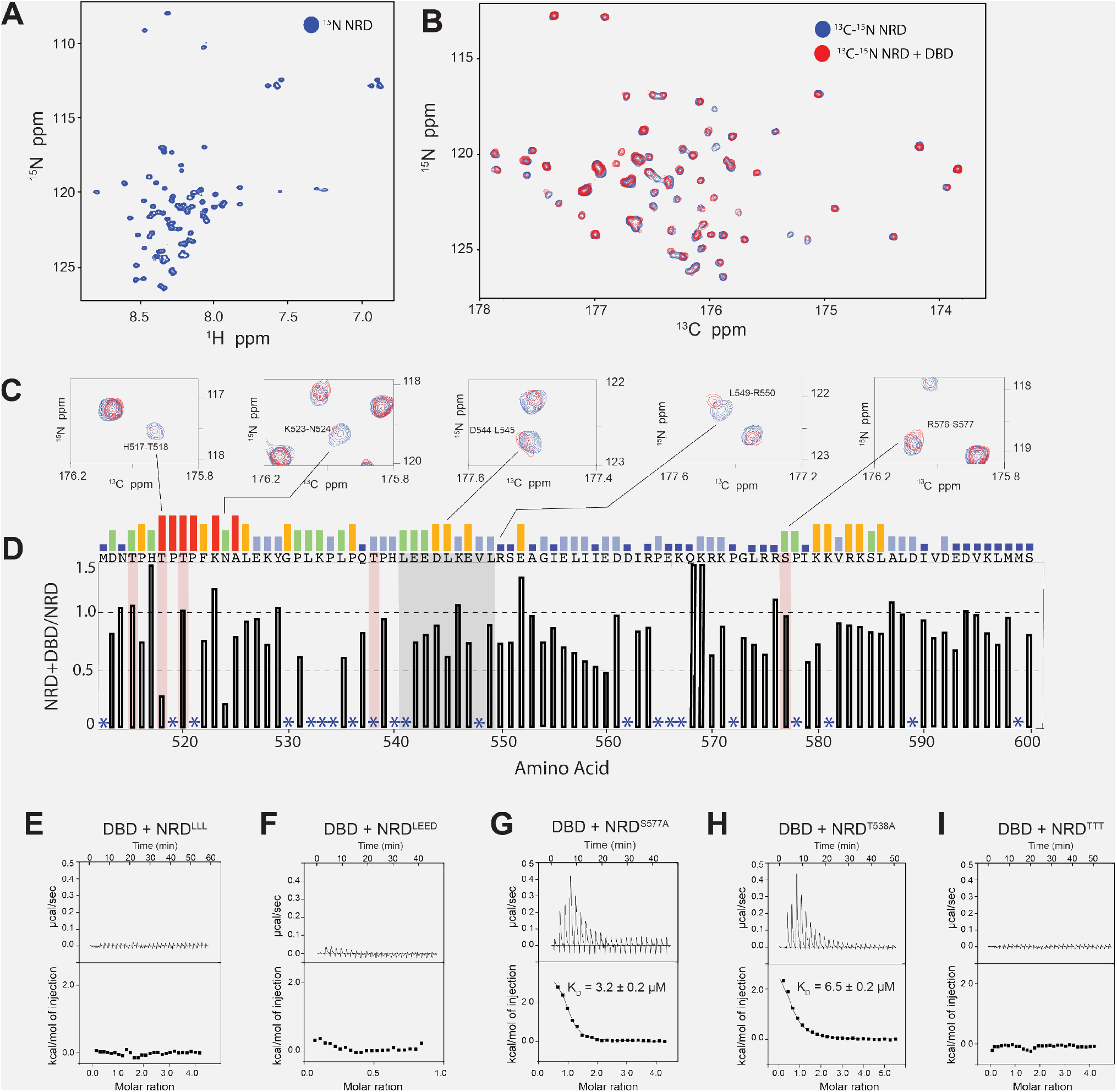
NMR spectroscopy maps potential NRD residues that interact with DBD. (A) ^1^H-^15^N HSQC spectrum of labeled NRD^510-600^ at 300 µM. (B) ^13^C-^15^N CON spectra of labeled NRD^510-600^ at 300 µM alone (blue) and with 600 µM unlabeled DBD (red). (C) Close-up views of exemplary assigned peaks in the ^13^C-^15^N CON spectra showing significant peak broadening. (D) Relative intensity of each amino acid plotted as the ratio of the intensity of NRD alone to the intensity of NRD + DBD. Asterisks mark amino acids that could not be assigned. Relative sequence conservation through Myb family members is displayed by the height of the bars above the primary sequence at the top of the graph. See Fig. S1 for the full NRD sequence alignment. (E-I) ITC binding measurements of DBD to mutant NRD^510-600^ constructs. LLL refers to L541, L545 and L549, LEED refers to L541, E542, E543, D544 and TTT refers to T515A, T518A, T520A.

We assigned the CON spectrum using standard backbone correlation experiments, and these assignments enabled identification of amino acid sequences in the NRD^510-600^ that are potentially critical for the interaction (Fig. 3C and 3D). A plot of peak intensity loss upon addition of DBD to the NRD^510-600^ sample shows that perturbations occurred at regions throughout NRD^510-600^ (Fig. 3D). We were particularly interested in the perturbations that clustered around residues 514-526 and 542-547 (Fig. 3D and Fig. S1). These clusters of residues show broadening, and the sequences are relatively well conserved. In addition, the sequence between 538-565 has helical propensity, and analysis suggest a hydrophobic surface containing several leucines that are conserved in B-Myb orthologs (Fig. S3). We surmised that if formed upon binding, such a helix would be a good candidate for facilitating interdomain interactions. To determine whether these regions are important for NRD-DBD association, we made two sets of alanine mutations in the most conserved residues found in these regions; we mutated together L541, E542, E543, D544 (NRD^LEED^) and together L541, L545 and L549 (NRD^LLL^). We expressed and purified the mutant NRD^510-600^ constructs and tested binding to DBD using ITC. We found that these mutations did not bind to DBD (Figs. 3E and 3F), supporting our NMR data that residues within 540-550 make critical contacts with the DBD.

There are five consensus Cdk2-CycA phosphorylation sites in NRD^510-600^ (T515, T518, T520, T538, and S577), and all of these phosphosites except T538 have been validated by two-dimensional tryptic peptide mapping and point mutagenesis (17,30,34,35). In our NMR spectra, we were unable to assign all the phosphorylation sites due to repetitive amino acid sequences, but we successfully assigned S577 and T518 (Fig. 3D). We observed a substantial change in intensity for the peak corresponding to T518 and for peaks corresponding to nearby residues (N514, T515, H517, T518) when DBD was added (Figs. 3C and 3D). In contrast, we observed minimal perturbations for the S577 peak and peaks corresponding to the residues around S577, which were reported to disrupt the NRD-DBD interaction when deleted (29). To probe the role of phosphorylation sites in the NRD-DBD interaction, we created three C-terminal NRD^510-600^ fragments with different phosphosites mutated to alanine (NRD^S577A^, NRD^T538A^, and NRD^TTT^, which contains T515A, T518A, T520A). We used these mutated and not phosphorylated NRDs in ITC experiments to detect binding affinities with DBD. We found that both NRD^S577A^ and NRD^T538A^ bound to DBD with K_D_ = 3.2 ± 0.2 µM and K_D_ = 6.5 ± 0.2 µM, respectively (Figs. 3G and 3H). However, NRD^TTT^ show no detectable binding, indicating that these threonines, when unphosphorylated, are important to interact with DBD (Fig. 3I).

### Phosphorylation of Cdk consensus sites in the conserved region of the NRD modulates the NRD-DBD interaction to regulate DNA binding

We next tested to what extent phosphorylation of Cdk sites in NRD^510-600^ influences NRD binding to the DBD and the inhibition of DBD-DNA binding. We phosphorylated purified NRD^510-600^ constructs with Cdk2-CycA and tested NRD-DBD affinities using ITC (Fig. 4). We verified phosphorylation of the WT NRD^510-600^ (B-Myb 510-600) on five sites with electrospray mass spectrometry (Fig. S2E). We observed no detectable binding towards DBD when WT NRD was phosphorylated (Fig. 4A). Similarly, we observed that phosphorylated NRD^S577A^ and NRD^T538A^ did not bind to DBD, indicating that phosphorylation of those specific sites is not required and that phosphorylation of T515A, T518A, T520A is sufficient to disrupt the interaction (Fig. 4B, 4C). Phosphorylation of NRD^TTT^ also resulted in no binding to DBD (Fig. 4D), although it was already established that these threonine residues were critical for the association when the protein is unphosphorylated (Fig. 3I).

**Figure 4.**
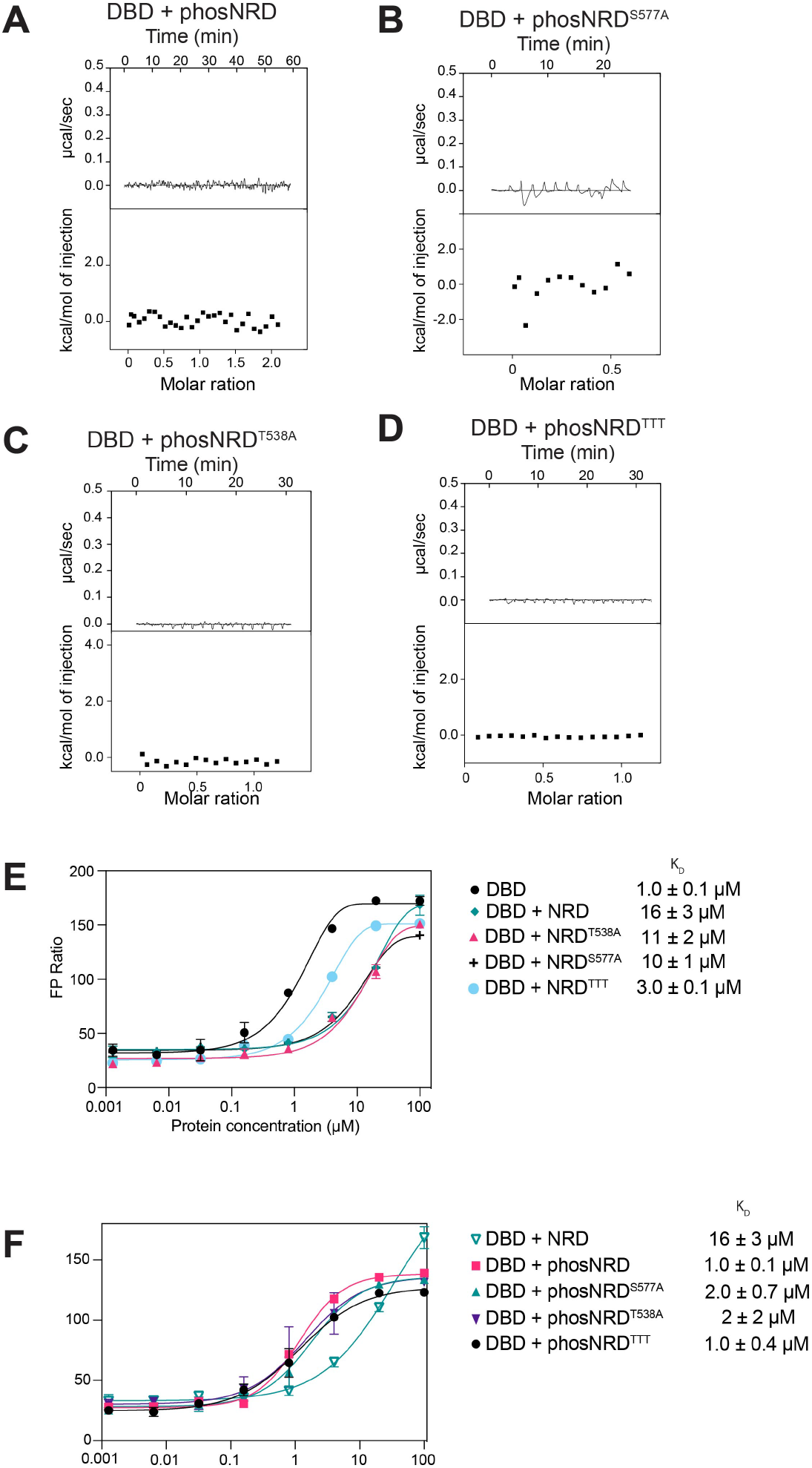
Phosphorylated NRD does not interact with DBD allowing DBD to interact with MBS DNA. (A-D) ITC measurements of DBD binding to the indicated NRD construct after phosphorylation by Cdk2-CyclinA. Phosphorylation of NRD was confirmed through electrospray mass spectrometry shown in Supplementary Fig. 1. (E-F) FP assay measurements of DBD binding to the TAMRA-labeled MBS probe incubated with the indicated NRD construct at 30 µM.

We performed the FP binding assay with fluorescently labeled MBS probe and added the various WT and phosphorylation-site mutant NRD^510-600^ constructs *in trans* (Figs. 4E and 4F). As previously shown in Fig. 1F, the DBD alone binds to the MBS probe with K_D_ of 1.1 ± 0.1 µM and when NRD is added *in trans* to DBD the affinity decreases to a K_D_ of 16 ± 3 µM. We found that, when unphosphorylated, the mutants NRD^S577A^ and NRD^T538A^ still inhibited DBD binding to the MBS probe. When NRD^TTT^ was added *in trans*, DBD binding affinity to the MBS probe was more weakly inhibited, consistent with our observation that T515, T518 and T520 are important for the interaction between the NRD and DBD that inhibits DBD binding to DNA (Fig. 4E). Phosphorylation of the NRD^S577A^ and NRD^T538A^ constructs with Cdk2-CycA abrogated their inhibitory effect on DNA binding, but phosphorylation of NRD^TTT^ resulted in DNA binding inhibition similar to the unphosphorylated mutant (Fig. 4F). Together these FP and ITC results are consistent with a model in which phosphorylation of T515, T518, T520 modulate the association of the NRD with the DBD in a manner that can regulate DNA binding affinity.

### Disruption of the NRD-DBD interaction increases the transactivation potential of B-Myb

To probe the functional significance of the NRD-DBD interaction in B-Myb-mediated transcriptional activation, we performed luciferase reporter assays in HCT116 cells (Fig. 5). Plasmids encoding wild-type and mutant B-Myb were transfected along with the pGL4.10 luciferase reporter plasmid containing an artificial promoter with three MBS consensus sequences. Such constructs have been previously utilized to detect B-Myb-dependent gene activation (36,37). As previously described, we observe a positive effect of B-Myb on the activity of the MBS promoter and a significant decrease of activation when the DBD is deleted. We tested mutation of phosphorylation sites in the NRD that we found to be important for NRD-DBD association in the NMR and ITC assays (Figs. 3 and 4). We found that B-Myb with mutation of the three NRD Cdk site threonines (T515/T518/T520) to either alanine or glutamate showed higher activity in the luciferase assay. Considering these mutations resulted in loss of NRD-DBD association, we propose that disruption of the repressive interaction leads to the observed more efficient B-Myb transactivation. We also tested two other mutations at Cdk sites in the NRD that were previously shown to regulate B-Myb by modulating the repressive activity of the NRD. However, in our assay, we found T538 mutation did not change B-Myb activity significantly from that of WT and the S577 mutation resulted in a subtle, albeit significant, reduction of activity.

**Figure 5.**
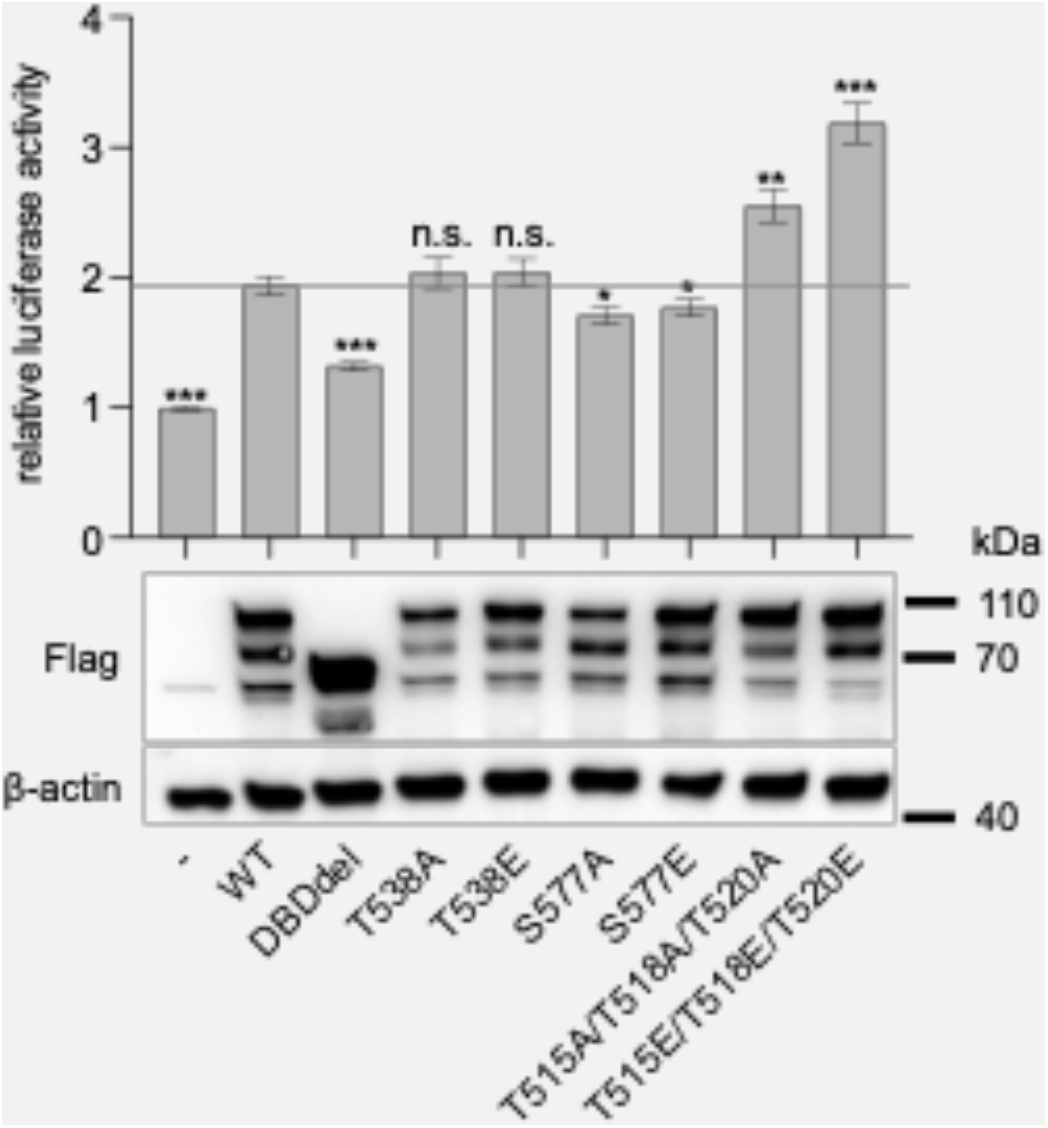
Disruption of a critical NRD-DBD interface hyper-activates B-myb. HCT116 cells were transfected with a luciferase reporter plasmid containing three Myb binding sites (MBS) upstream of a minimal promoter together with plasmids expressing Flag-tagged wild-type B-Myb or the indicated mutants. Mean values ± SD of four biological replicates are given, and significances were calculated by the Students paired T-Test (*p≤0.05, **p≤0.01, ***p≤0.001 compared with wild-type (WT) B-Myb). Expression levels of B-MYB variants in the luciferase assay samples were analyzed by SDS-PAGE/Western blot.

## Discussion

Our data show that B-Myb binding to an MBS DNA sequence is inhibited by the intramolecular association between the DBD and the NRD region between 510-600 (Fig. 6). This inhibited conformation is regulated by Cdk2-CycA dependent phosphorylation of T515, T518, and T520, which disrupts the interdomain interaction between the NRD and DBD and permits stronger DNA association. Our mechanistic findings are generally consistent with a number of studies demonstrating, primarily using cell-based reporter assays, that B-Myb transactivation of MBS promoters is autoinhibited by the NRD and increased by co-transfection with CycA (17,25-27,34,35,38). Moreover, our biochemical data that Cdk phosphorylation specifically modulates DNA binding offers mechanistic explanation for previous observations that B-Myb phosphorylation and localization to target promoters are coincident (24).

**Figure 6.**
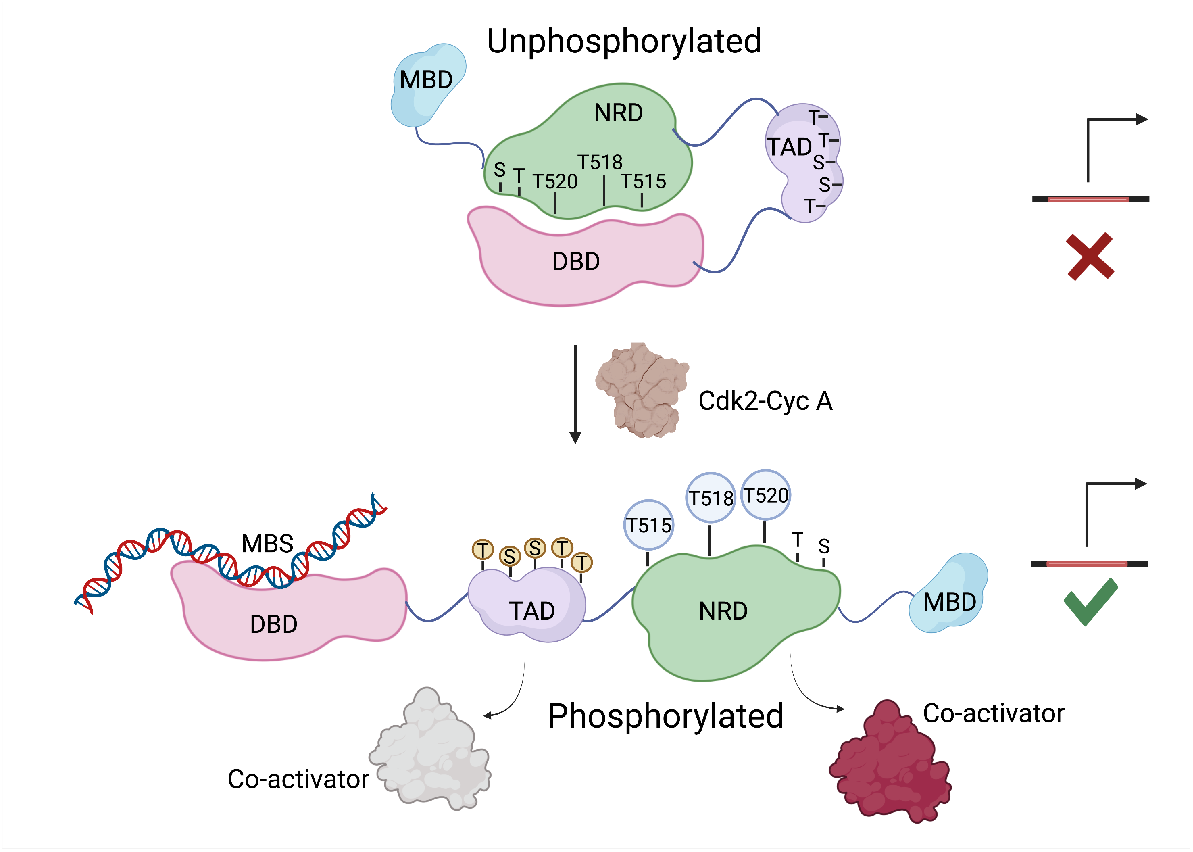
Structural model for B-Myb autoinhibition and activation upon Cdk2 phosphorylation.

We note several differences between our findings here monitoring the behavior of purified proteins and previous results from cell-based assays. For example, previous studies determined that C-terminal truncations starting from D561 cause the strongest hyperactivity of B-Myb towards a promoter containing Myb binding sites (17,27). Co-transfection with Cdk2-CycA further stimulated the activity of the truncations; however, the activity of full-length B-MYB was much further increased by Cdk2-CycA overexpression (27). More recently, it was reported that the B-Myb DBD interacts with a region in the NRD between 560-589 and Cdk mediated phosphorylation of the residue S577 relieves this inhibition (31). In contrast, our NMR and mutagenesis data from biochemical and reporter assays implicate sequences in the NRD that are N-terminal to D561 as those making primary contact with and regulating the DBD, including sequences around the T515, T518 and T520 phosphorylation sites and the amino acid stretch from L541 to L549 (Fig. 3). Our studies did not find S577 to be involved in regulating the NRD-DBD association or S577E to have a positive effect on MBS promoter activity. Rather, we found the more conserved T515/T518/T520 as important phosphorylation sites that regulate MBS-dependent activity. In contrast, other studies using reporter-based cell assays found that point mutations of T518/T518/T520 inhibit MBS transactivation (26,30,39); however, it should be noted that other phosphorylation sites were mutated in addition to these sites and may function through independent mechanisms. As an overarching explanation to differences between previous studies and our results, which specifically focus on DNA binding, we speculate that other protein interactions or additional posttranslational modifications also account for the importance of the NRD and its phosphorylation for B-Myb activity and regulation.

Our results demonstrate how intrinsically disordered regions (IDRs) in TFs can regulate TF interactions and how this regulation can be modulated through posttranslational modifications. Other examples of IDRs specifically influencing TF binding to DNA include p53, PU1, ETS1, and TFB2M (40-43). Like B-Myb, other proteins that control the cell cycle are typically phosphorylated at multiple sites in their IDRs by Cdks (44-47). Multi-site phosphorylation of TFs like B-Myb and their regulators mediates unique functions through significant structural changes in interdomain interactions. Phosphorylation-dependent structural changes alter conformations from disorder to order to facilitate or inhibit interactions. For example, phosphorylation of the mitotic transcription factor FoxM1 by Cdk2-CycA and Plk1 switches the protein conformation from an inactive to an active state by inhibiting intra-molecular interactions (45). Our observation that B-Myb inhibition of NRD-DBD released upon Cdk mediated phosphorylation aligns with this common theme of regulation in cell-cycle transcription factors through control of interdomain interactions and structural transitions that promote or reduce structural disorder.

Phosphorylation has been shown to regulate activation of the other Myb family transcription factors. c-Myb gets phosphorylated by several kinases other than Cdk2, including CK2, protein kinase A (PKA), and mitogen activated protein kinase (MAPK) (48-50). There is evidence that the NRDs of c-Myb and A-Myb regulate DNA binding as well; for example, C-terminal truncations show increased DNA binding and activation of the Myb dependent reporter promoter *mim-1* (18,51). The NRD of c-Myb interacts with its DBD though a conserved EVES motif; however, this inhibitory interaction is stimulated by phosphorylation of S581 (the S in EVES) (48,52). In fact, phosphorylation-dependent rescue of the negative regulation of NRD is not reported for either A-Myb or c-Myb. Therefore, even though Myb proteins have an evolutionarily conserved DBD and recognize the same DNA motif, different phosphorylation patterns might follow unique functions in regard to their tissue-specific functions.

Our study and most previous studies of B-Myb activation consider B-Myb as a site-specific transcription factor that transactivates from MBS sites, and thus gene expression assays primarily monitor artificial Myb-responsive reporter genes (17,35,36) or type I collagen promoter activity (25). Some studies of B-Myb in the context of activating cell-cycle transcription have led to a model for B-Myb function in which it acts independently of DNA binding to the MBS site. In particular, the MBS site is not essential to activate genes that show cell-cycle dependent expression and contain the CHR site that is bound by MuvB (53). In contrast, the association of B-Myb with MuvB is critical for gene activation and proper localization of B-Myb to promoters (23,24,54). In one remarkable example in flies, expression of the MuvB-binding domain is necessary and sufficient to restore the activity of the PLK1 promoter after loss of B-Myb, suggesting that the DBD is not necessary for B-Myb function in activating some genes (22). It may be then that activation of B-Myb’s ability to bind DNA by Cdk mediated phosphorylation is a particular mechanism for MBS promoters, while activation of CHR promoters may entail other additional mechanisms. Nevertheless, a slow migrating phosphorylated form of B-Myb is abundantly found in S-phase of the cell cycle which coincides with B-Myb expression and localization to cell-cycle dependent promoters (29,35,55). Thus, it is possible that phosphorylation might be enhancing other protein-protein interactions like FoxM1 that are specifically necessary for cell-cycle dependent gene activation (24). Further studies on B-Myb will shed light on how it activates cell-cycle dependent genes at the CHR promoters, and it will be interesting to further identify relevant MBS-dependent genes that respond to cell-cycle control of B-Myb activity by Cdk2-CycA.

## Materials and Methods

### Recombinant protein expression and purification

The human B-Myb full-length protein and B-Myb^34-600^ constructs were expressed in Sf9 cells with a cleavable N-terminal Strep tag using the FastBac expression system. Cells were harvested and lysed in a buffer containing 300 mM NaCl, 50 mM Tris, 1 mM DTT, 10% Glycerol v/v, Sigma Protease Inhibitor (P8340) and 1 mM phenylmethylsulfonyl fluoride (pH 8.0). Protein was purified with StrepTactin™ Sepharose™ High Performance resin (Cytvia) equilibrated in lysis buffer. The lysed cells were clarified by centrifugation at 19,000 rpm for 45 minutes at 4^°^ C. The cleared lysate was incubated with the resin was washed with a buffer containing 300 mM NaCl, 50 mM Tris, 1 mM DTT, 10% glycerol v/v (pH 8.0) for 1 hour at 4^°^ C and washed to remove unspecific proteins. The protein was then eluted in 300 mM NaCl, 50 mM Tris, 5 mM desthiobiotin, 10% glycerol v/v, 1 mM DTT (pH 8.0). Protein was dialyzed into storage buffer (200 mM NaCl, 50 mM Tris, 1 mM BME, 10% glycerol v/v (pH 8.0)) and stored at −80 °C.

The human B-Myb truncated constructs (DBD, TAD, NRD, B-Myb^34-370^) were expressed in *E. coli* from an engineered pGEX plasmid with an N-terminal GST tag and a TEV protease cleavage site. Proteins were expressed overnight by inducing with 1 mM IPTG at 19 ^°^C. All proteins were lysed in a buffer containing 200 mM NaCl, 40 mm Tris, 5 mM DTT, 1 mM phenylmethylsulfonyl fluoride (pH 8.0). The lysed cells were clarified by centrifugation at 19,000 rpm for 45 min at 4^°^ C. Protein lysates were allowed to bind to equilibrated Glutathione Sepharose resin (Cytvia) for 30 minutes and washed to remove unspecific proteins. The proteins were eluted with a buffer containing 200 mM NaCl, 40 mm Tris, 5 mM DTT, 10 mM L-Glutathione reduced (pH 8.0). Eluted proteins were further purified using Q-sepharose and cleaved with TEV protease at 4^°^ C overnight. Proteins were then passed through Glutathione Sepharose resin to remove the free GST and concentrated to run through Superdex-75 (GE Healthcare) into 200 mM NaCl, 25 mM Tris, 1 mM DTT (pH 8.0). Cdk2-CycA and Plk1 kinase domains were expressed and purified as previously described (45).

To generate phosphorylated protein reagents, kinase reactions were performed similar to as previously described (45). B-Myb protein constructs following final purification were incubated with 10 mM ATP, 50 mM MgCl_2_, and 20% by mass of either Cdk2-CycA, Plk1 kinase domain, or both Plk1 and Cdk2-CycA, overnight at 4°C. The kinase reaction was concentrated and run over Superdex-75 (GE Healthcare) to remove kinases and ATP and phosphorylation of the proteins were confirmed by electrospray mass spectrometry using a Sciex X500B QTOF system.

### Fluorescence polarization assay

Dissociation constants for direct binding between DBD and MBS DNA sequence were determined using increasing amounts of DBD titrated into 20 nM of fluorescently labeled MBS DNA probe. For DBD + NRD assays, DBD and NRD were incubated for 30 minutes on ice before titrating the labeled MBS probe in a buffer containing 150 mM NaCl, 25 mM Tris, 1 mM DTT, 0.1% Tween20 (pH 8). FP measurements were measured on a Perkin-Elmer EnVision 2103 Multilabel plate reader with excitation at 523 nm and emission at 580 nm. The dissociation constants (K_D_) were calculated by fitting milipolarization (mP) values of three technical replicates against concentration using a one site binding model in GraphPad Prism 8.

### Isothermal titration calorimetry

Dissociation constants (K_D_) for DBD and NRD interactions were measured using ITC with a MicroCal VP-ITC system. All proteins were concentrated as needed and dialyzed into a buffer containing 150 mM NaCl, 20 mM Tris, 1 mM BME (pH 8). DBD (500 µM) was titrated into NRD (50 µM) at 19 ^°^ C. The dissociation constant of NRD mutants and phosphorylated NRDs were determined similarly. K_D_s are the average fits from three technical replicates analyzed using the Origin ITC software package with the standard deviation reported as error.

### NMR spectroscopy

The HSQC and CON spectra for DBD and NRD interaction studies in Fig. 3 were collected at 25 ^°^ C on a Bruker Avance III HD 800-MHz spectrometer equipped with cryogenically cooled probes. The sample contained ^13^C-^15^N-labeled NRD^510-600^ at 300 µM in a buffer containing 20 mM sodium phosphate pH 8.0, 100 mM NaCl, 1 mM DTT and 5% (v/v) D_2_O. The backbone assignment of the NRD was accomplished using standard NH-edited triple-resonance experiments [HNCO, HNCACB, CBCA(CO)NH, C(CO)NH] supplied by Varian/Agilent (33). The NH-edited experiments were collected on a Varian/Agilent INOVA 600 MHz NMR equipped with cryogenically cooled probes. The experiments for backbone assignments were collected at pH 6.0 (otherwise same buffer) due to favorable chemical exchange and assignments were transferred to pH 8.0 CON spectra through pH titrating. All spectra were processed using NMRPipe and analyzed and assigned using Sparky (60,61).

### Cell culture, luciferase assays, and Western blot

HCT116 colon carcinoma cells were grown in Dulbecco’s modified Eagle’s medium (Gibco, high glucose, GlutaMAX Supplement, pyruvate) supplemented with 10 % fetal bovine serum (Corning, Regular Fetal Bovine Serum) and penicillin/streptomycin (Gibco). Cells were maintained at 37° C and 5 % CO2.

The 3xMBS luciferase reporter construct was created by inserting a double-stranded oligonucleotide containing three copies of a high affinity B-MYB binding site (TAACGGTG) (1-4) upstream of the herpes simplex thymidine kinase (TK) minimal promoter (5-TTA**TAACGGTC**TTAA**TAACGGTC**TTAA**TAACGGTC**TTTTAGC*TTCGCATATTAAGGTGACGCGT GTGGCCTCGAACACCGAGCGACCCTGCAGCGACCCGCTTAA*-3; MBSs in bold, minimal TK promoter in italics) into the KpnI and NcoI sites of the pGL4.10[luc2] vector (Promega). The ORF of human MYBL2/B-Myb isoform 1 (NM_002466.4) was cloned into pcDNA3.1+ (ThermoFisher Scientific) and fused with an N-terminal Flag tag. Point mutations were introduced following the QuikChange site-directed mutagenesis protocol, and the DBD was deleted following the NEB Q5 protocol.

Stimulation of the 3xMBS promoter activity was analyzed by luciferase reporter assays with extracts of transfected HCT116 cells. 30,000 cells per 48 well were plated and transfected with 1 μl PEI (Polysciences, PEI 25K), 75 ng of promoter reporter plasmids (3xMBS-pGL4.10 or pGl4.10 empty vector) along with 100 ng of pcDNA3.1 plasmids expressing Flag-B-Myb (wild-type or mutants) and 25 ng renilla luciferase plasmid (pGL4.70). Cells were lysed 48h after transfection, and luciferase activity was measured with the Dual-Luciferase Reporter Assay System (Promega) following the manufacturer’s recommendations on an EnVision 2105 plate reader (PerkinElmer). Relative promoter activities of the 3xMBS-pGL4.10 reporter after expression of wild-type or mutant B-MYB were calculated by normalizing to renilla luciferase activity and to the activity of the pGL4.10 empty vector co-transfected with the respective B-MYB constructs.

Expression levels of wild-type and mutant B-MYB were analyzed by loading 10 µg of the remaining luciferase assay lysates onto a 10% SDS gel followed by Western blotting. Flag-B-MYB was detected with the Anti-OctA-Probe antibody (Santa Cruz Biotech, sc-166355 HRP, dilution 1:2,000), and β-actin was probed with the Direct-Blot HRP anti-β-actin antibody (BioLegend, clone W16197A, Cat. # 664804, dilution 1:10,000).

## Supplementary Figures

**Supplementary Figure 1.**
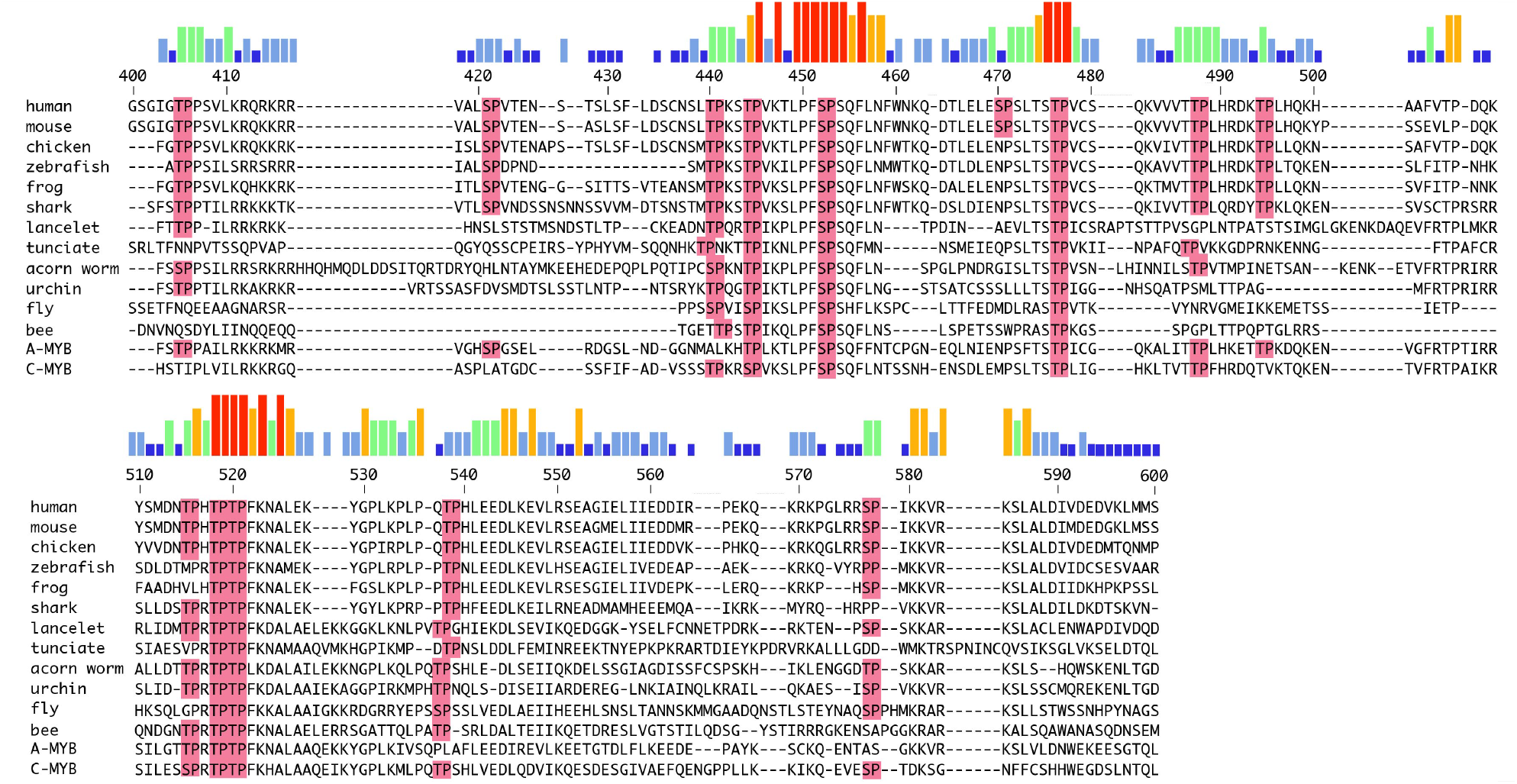
Sequence alignment of the B-Myb NRD. Sequences for B-Myb orthologs are shown along with the sequences of human A-MYB and c-MYB. Minimal Cdk sites consensus sites (S/TP) are highlighted in pink.

**Supplementary Figure 2.**
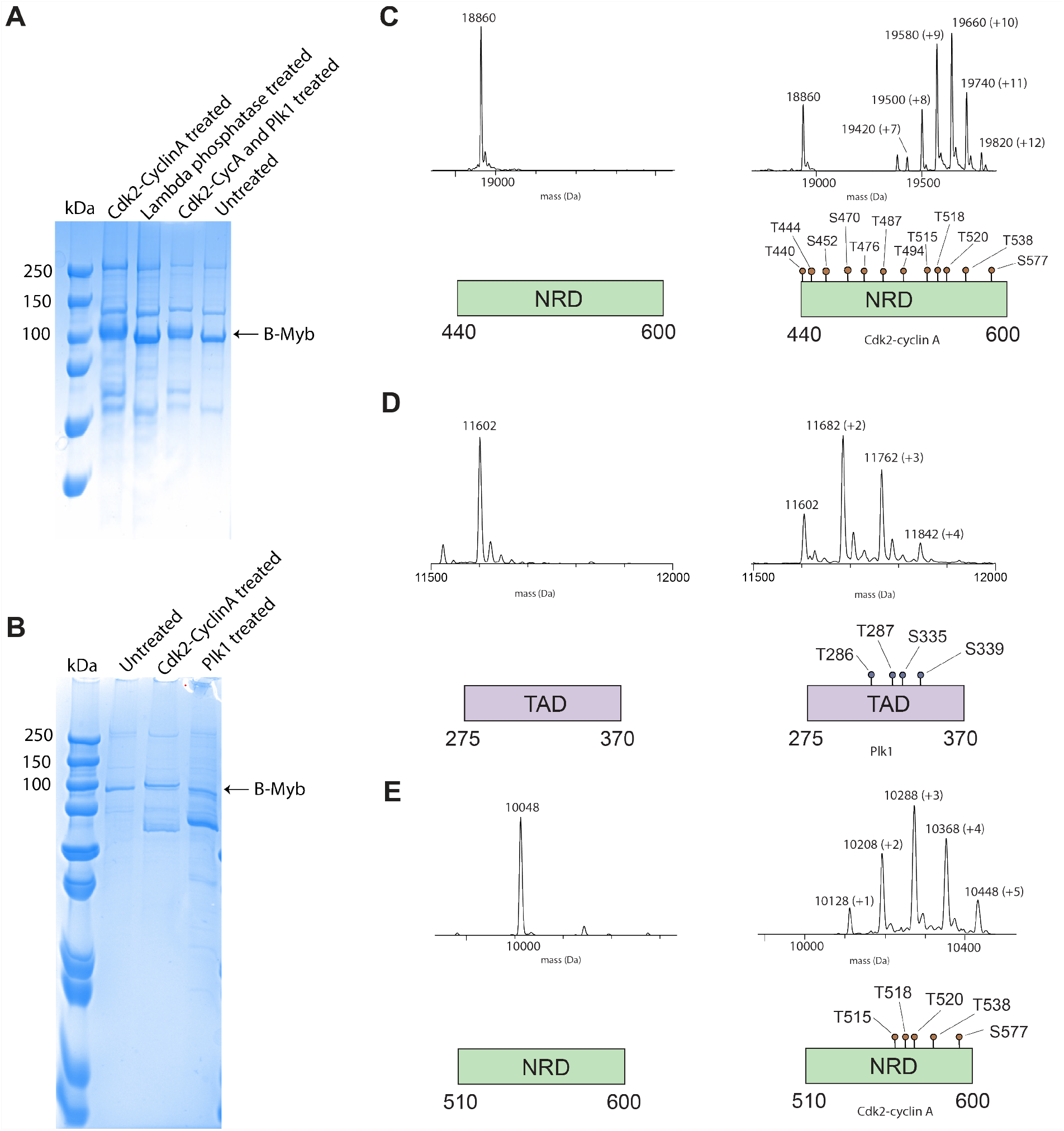
Confirmation of recombinant protein phosphorylation. (A-B) PhosTag gels following purification of full-length B-Myb expressed in Sf9 cells and treatment with the indicated enzyme(s). The B-Myb band appears with similar mobility in the phosphatase treated and untreated samples, suggesting that untreated purified protein is not extensively phosphorylated. Phosphorylation with Cdk2-CycA produces a noticeable shift, whereas the shift with Plk1 treatment is not detectable when using Plk1 alone or together with Cdk2-CycA. However, we are confident in the activity of our Plk1 enzyme preparation considering our mass spectrometry results detecting TAD phosphorylation in panel D. (C) B-Myb NRD^440-600^ unphosphorylated (left) and phosphorylated (right) by Cdk2-Cyc A detected by mass spectrometry. In this construct there are 3 strong Cdk consensus sites ({S/T}Px{K/R}) at T444, T520 and S577 and 9 other weak consensus sites ({S/T}P). Peaks are labeled with the final mass and the number of phosphoryl groups added in parenthesis (+80 Da). (D) B-Myb TAD unphosphorylated (left) and phosphorylated (right) by Plk1. This construct contains 4 Plk1 consensus sites ({D/N/E/Q}xS!) as depicted in the schematic diagram. (E) B-Myb NRD^510-600^ unphosphorylated (left) and phosphorylated (right) by Cdk2-Cyc A detected by mass spectrometry.

**Supplementary Figure 3.**
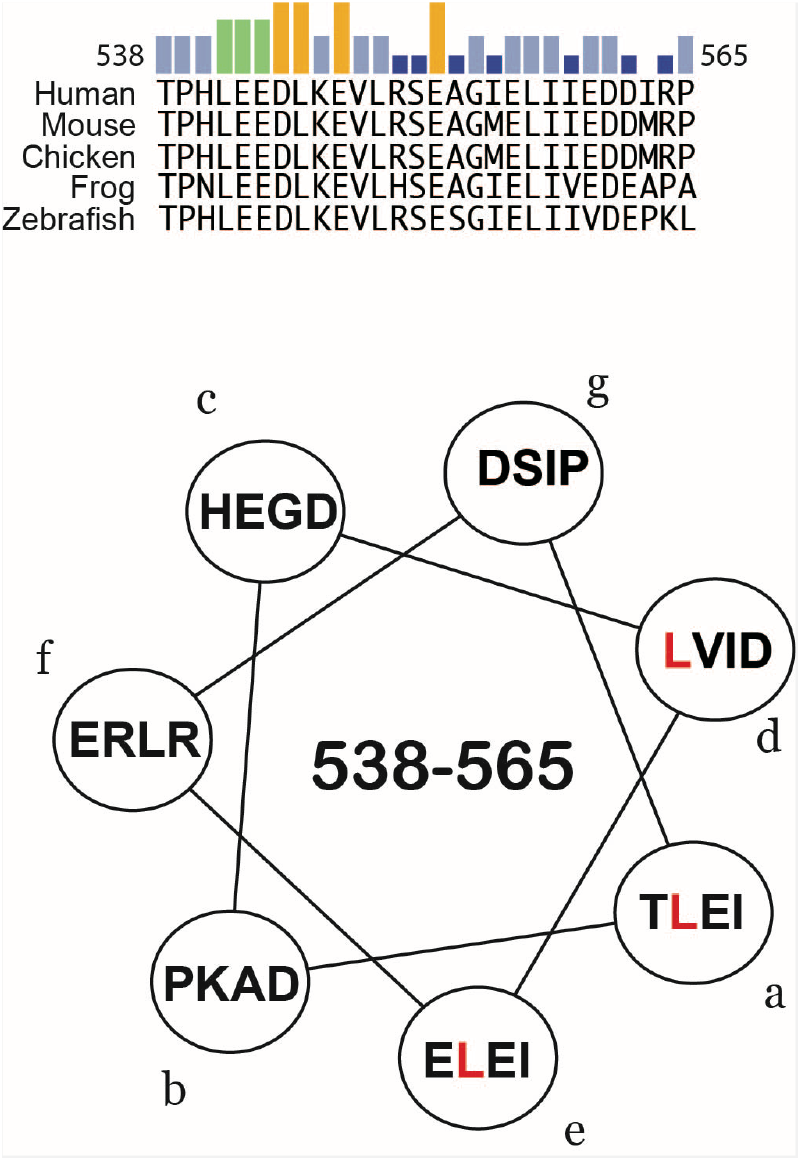
Rational for probing L541/L545/L549 NRD mutant. (Top) Sequences for five B-Myb orthologs. The bars on top of the sequence reflect conservation from the larger sequence alignment shown in Fig. S1. (Bottom) Helical wheel projection beginning with T538 in the “a” position. The projection suggests that this sequence has the potential to form an amphipathic helix with the three leucines (colored in red) positioned along one face.

## References

1. Bergholtz, S., Andersen, T. O., Andersson, K. B., Borrebaek, J., Luscher, B., and Gabrielsen, O. S. (2001) The highly conserved DNA-binding domains of A-, B-and c-Myb differ with respect to DNA-binding, phosphorylation and redox properties. Nucleic Acids Res 29, 3546–3556

2. Carr, M. D., Wollborn, U., McIntosh, P. B., Frenkiel, T. A., McCormick, J. E., Bauer, C. J., Klempnauer, K. H., and Feeney, J. (1996) Structure of the B-Myb DNA-binding domain in solution and evidence for multiple conformations in the region of repeat-2 involved in DNA binding: implications for sequence-specific DNA binding by Myb proteins. Eur J Biochem 235, 721–735

3. Graf, T. (1992) Myb: a transcriptional activator linking proliferation and differentiation in hematopoietic cells. Curr Opin Genet Dev 2, 249–255

4. Lipsick, J. S. (1996) One billion years of Myb. Oncogene 13, 223–235

5. Nitta, K. R., Jolma, A., Yin, Y., Morgunova, E., Kivioja, T., Akhtar, J., Hens, K., Toivonen, J., Deplancke, B., Furlong, E. E., and Taipale, J. (2015) Conservation of transcription factor binding specificities across 600 million years of bilateria evolution. Elife 4

6. Biedenkapp, H., Borgmeyer, U., Sippel, A. E., and Klempnauer, K. H. (1988) Viral myb oncogene encodes a sequence-specific DNA-binding activity. Nature 335, 835–837

7. Ogata, K., Morikawa, S., Nakamura, H., Sekikawa, A., Inoue, T., Kanai, H., Sarai, A., Ishii, S., and Nishimura, Y. (1994) Solution structure of a specific DNA complex of the Myb DNA-binding domain with cooperative recognition helices. Cell 79, 639–648

8. Oh, I. H., and Reddy, E. P. (1999) The myb gene family in cell growth, differentiation and apoptosis. Oncogene 18, 3017–3033

9. Mucenski, M. L., McLain, K., Kier, A. B., Swerdlow, S. H., Schreiner, C. M., Miller, T. A., Pietryga, D. W., Scott, W. J., Jr., and Potter, S. S. (1991) A functional c-myb gene is required for normal murine fetal hepatic hematopoiesis. Cell 65, 677–689

10. Toscani, A., Mettus, R. V., Coupland, R., Simpkins, H., Litvin, J., Orth, J., Hatton, K. S., and Reddy, E. P. (1997) Arrest of spermatogenesis and defective breast development in mice lacking A-myb. Nature 386, 713–717

11. Sala, A. (2005) B-MYB, a transcription factor implicated in regulating cell cycle, apoptosis and cancer. Eur J Cancer 41, 2479–2484

12. Bayley, R., Ward, C., and Garcia, P. (2020) MYBL2 amplification in breast cancer: Molecular mechanisms and therapeutic potential. Biochim Biophys Acta Rev Cancer 1874, 188407

13. Lin, D., Fiscella, M., O′Connor, P. M., Jackman, J., Chen, M., Luo, L. L., Sala, A., Travali, S., Appella, E., and Mercer, W. E. (1994) Constitutive expression of B-myb can bypass p53-induced Waf1/Cip1-mediated G1 arrest. Proc Natl Acad Sci U S A 91, 10079–10083

14. Qin, H., Li, Y., Zhang, H., Wang, F., He, H., Bai, X., and Li, S. (2019) Prognostic implications and oncogenic roles of MYBL2 protein expression in esophageal squamous-cell carcinoma. Onco Targets Ther 12, 1917–1927

15. Sun, C., Li, H., Mills, R. E., and Guan, Y. (2019) Prognostic model for multiple myeloma progression integrating gene expression and clinical features. Gigascience 8

16. Dubendorff, J. W., Whittaker, L. J., Eltman, J. T., and Lipsick, J. S. (1992) Carboxy-terminal elements of c-Myb negatively regulate transcriptional activation in cis and in trans. Genes Dev 6, 2524–2535

17. Lane, S., Farlie, P., and Watson, R. (1997) B-Myb function can be markedly enhanced by cyclin A-dependent kinase and protein truncation. Oncogene 14, 2445–2453

18. Takahashi, T., Nakagoshi, H., Sarai, A., Nomura, N., Yamamoto, T., and Ishii, S. (1995) Human A-myb gene encodes a transcriptional activator containing the negative regulatory domains. FEBS Lett 358, 89–96

19. Brayer, K. J., Frerich, C. A., Kang, H., and Ness, S. A. (2016) Recurrent Fusions in MYB and MYBL1 Define a Common, Transcription Factor-Driven Oncogenic Pathway in Salivary Gland Adenoid Cystic Carcinoma. Cancer Discov 6, 176–187

20. Gonda, T. J., Buckmaster, C., and Ramsay, R. G. (1989) Activation of c-myb by carboxy-terminal truncation: relationship to transformation of murine haemopoietic cells in vitro. EMBO J 8, 1777–1783

21. Lipsick, J. S., and Wang, D. M. (1999) Transformation by v-Myb. Oncogene 18, 3047–3055

22. Andrejka, L., Wen, H., Ashton, J., Grant, M., Iori, K., Wang, A., Manak, J. R., and Lipsick, J. S. (2011) Animal-specific C-terminal domain links myeloblastosis oncoprotein (Myb) to an ancient repressor complex. Proc Natl Acad Sci U S A 108, 17438–17443

23. Guiley, K. Z., Iness, A. N., Saini, S., Tripathi, S., Lipsick, J. S., Litovchick, L., and Rubin, S. M. (2018) Structural mechanism of Myb-MuvB assembly. Proc Natl Acad Sci U S A 115, 10016–10021

24. Sadasivam, S., Duan, S., and DeCaprio, J. A. (2012) The MuvB complex sequentially recruits B-Myb and FoxM1 to promote mitotic gene expression. Genes Dev 26, 474–489

25. Petrovas, C., Jeay, S., Lewis, R. E., and Sonenshein, G. E. (2003) B-Myb repressor function is regulated by cyclin A phosphorylation and sequences within the C-terminal domain. Oncogene 22, 2011–2020

26. Ziebold, U., Bartsch, O., Marais, R., Ferrari, S., and Klempnauer, K. H. (1997) Phosphorylation and activation of B-Myb by cyclin A-Cdk2. Curr Biol 7, 253–260

27. Bessa, M., Saville, M. K., and Watson, R. J. (2001) Inhibition of cyclin A/Cdk2 phosphorylation impairs B-Myb transactivation function without affecting interactions with DNA or the CBP coactivator. Oncogene 20, 3376–3386

28. Tashiro, S., Takemoto, Y., Handa, H., and Ishii, S. (1995) Cell type-specific trans-activation by the B-myb gene product: requirement of the putative cofactor binding to the C-terminal conserved domain. Oncogene 10, 1699–1707

29. Werwein, E., Cibis, H., Hess, D., and Klempnauer, K. H. (2019) Activation of the oncogenic transcription factor B-Myb via multisite phosphorylation and prolyl cis/trans isomerization. Nucleic Acids Res 47, 103–121

30. Johnson, T. K., Schweppe, R. E., Septer, J., and Lewis, R. E. (1999) Phosphorylation of B-Myb regulates its transactivation potential and DNA binding. J Biol Chem 274, 36741–36749

31. Werwein, E., Biyanee, A., and Klempnauer, K. H. (2020) Intramolecular interaction of B-MYB is regulated through Ser-577 phosphorylation. FEBS Lett 594, 4266–4279

32. Charrasse, S., Carena, I., Brondani, V., Klempnauer, K. H., and Ferrari, S. (2000) Degradation of B-Myb by ubiquitin-mediated proteolysis: involvement of the Cdc34-SCF(p45Skp2) pathway. Oncogene 19, 2986–2995

33. Bastidas, M., Gibbs, E. B., Sahu, D., and Showalter, S. A. (2015) A primer for carbon-detected NMR applications to intrinsically disordered proteins in solution. Concepts in Magnetic Resonance Part A 44, 54–66

34. Sala, A., Kundu, M., Casella, I., Engelhard, A., Calabretta, B., Grasso, L., Paggi, M. G., Giordano, A., Watson, R. J., Khalili, K., and Peschle, C. (1997) Activation of human B-MYB by cyclins. Proc Natl Acad Sci U S A 94, 532–536

35. Saville, M. K., and Watson, R. J. (1998) The cell-cycle regulated transcription factor B-Myb is phosphorylated by cyclin A/Cdk2 at sites that enhance its transactivation properties. Oncogene 17, 2679–2689

36. Ness, S. A., Marknell, A., and Graf, T. (1989) The v-myb oncogene product binds to and activates the promyelocyte-specific mim-1 gene. Cell 59, 1115–1125

37. Seong, H. A., Kim, K. T., and Ha, H. (2003) Enhancement of B-MYB transcriptional activity by ZPR9, a novel zinc finger protein. J Biol Chem 278, 9655–9662

38. Ansieau, S., Kowenz-Leutz, E., Dechend, R., and Leutz, A. (1997) B-Myb, a repressed trans-activating protein. J Mol Med (Berl) 75, 815–819

39. Bartsch, O., Horstmann, S., Toprak, K., Klempnauer, K. H., and Ferrari, S. (1999) Identification of cyclin A/Cdk2 phosphorylation sites in B-Myb. Eur J Biochem 260, 384–391

40. Sun, X., Dyson, H. J., and Wright, P. E. (2021) A phosphorylation-dependent switch in the disordered p53 transactivation domain regulates DNA binding. Proc Natl Acad Sci U S A 118

41. Basu, U., Mishra, N., Farooqui, M., Shen, J., Johnson, L. C., and Patel, S. S. (2020) The C-terminal tails of the mitochondrial transcription factors Mtf1 and TFB2M are part of an autoinhibitory mechanism that regulates DNA binding. J Biol Chem 295, 6823–6830

42. Pufall, M. A., Lee, G. M., Nelson, M. L., Kang, H. S., Velyvis, A., Kay, L. E., McIntosh, L. P., and Graves, B. J. (2005) Variable control of Ets-1 DNA binding by multiple phosphates in an unstructured region. Science 309, 142–145

43. Xhani, S., Lee, S., Kim, H. M., Wang, S., Esaki, S., Ha, V. L. T., Khanezarrin, M., Fernandez, G. L., Albrecht, A. V., Aramini, J. M., Germann, M. W., and Poon, G. M. K. (2020) Intrinsic disorder controls two functionally distinct dimers of the master transcription factor PU.1. Sci Adv 6, eaay3178

44. Fu, Z., Malureanu, L., Huang, J., Wang, W., Li, H., van Deursen, J. M., Tindall, D. J., and Chen, J. (2008) Plk1-dependent phosphorylation of FoxM1 regulates a transcriptional programme required for mitotic progression. Nat Cell Biol 10, 1076–1082

45. Marceau, A. H., Brison, C. M., Nerli, S., Arsenault, H. E., McShan, A. C., Chen, E., Lee, H. W., Benanti, J. A., Sgourakis, N. G., and Rubin, S. M. (2019) An order-to-disorder structural switch activates the FoxM1 transcription factor. Elife 8

46. Rubin, S. M. (2013) Deciphering the retinoblastoma protein phosphorylation code. Trends Biochem Sci 38, 12–19

47. Xu, M., Sheppard, K. A., Peng, C. Y., Yee, A. S., and Piwnica-Worms, H. (1994) Cyclin A/CDK2 binds directly to E2F-1 and inhibits the DNA-binding activity of E2F-1/DP-1 by phosphorylation. Mol Cell Biol 14, 8420–8431

48. Aziz, N., Miglarese, M. R., Hendrickson, R. C., Shabanowitz, J., Sturgill, T. W., Hunt, D. F., and Bender, T. P. (1995) Modulation of c-Myb-induced transcription activation by a phosphorylation site near the negative regulatory domain. Proc Natl Acad Sci U S A 92, 6429–6433

49. Luscher, B., Christenson, E., Litchfield, D. W., Krebs, E. G., and Eisenman, R. N. (1990) Myb DNA binding inhibited by phosphorylation at a site deleted during oncogenic activation. Nature 344, 517–522

50. Ramsay, R. G., Morrice, N., Van Eeden, P., Kanagasundaram, V., Nomura, T., De Blaquiere, J., Ishii, S., and Wettenhall, R. (1995) Regulation of c-Myb through protein phosphorylation and leucine zipper interactions. Oncogene 11, 2113–2120

51. Hu, Y. L., Ramsay, R. G., Kanei-Ishii, C., Ishii, S., and Gonda, T. J. (1991) Transformation by carboxyl-deleted Myb reflects increased transactivating capacity and disruption of a negative regulatory domain. Oncogene 6, 1549–1553

52. Dash, A. B., Orrico, F. C., and Ness, S. A. (1996) The EVES motif mediates both intermolecular and intramolecular regulation of c-Myb. Genes Dev 10, 1858–1869

53. Muller, G. A., Wintsche, A., Stangner, K., Prohaska, S. J., Stadler, P. F., and Engeland, K. (2014) The CHR site: definition and genome-wide identification of a cell cycle transcriptional element. Nucleic Acids Res 42, 10331–10350

54. Down, C. F., Millour, J., Lam, E. W., and Watson, R. J. (2012) Binding of FoxM1 to G2/M gene promoters is dependent upon B-Myb. Biochim Biophys Acta 1819, 855–862

55. Robinson, C., Light, Y., Groves, R., Mann, D., Marias, R., and Watson, R. (1996) Cell-cycle regulation of B-Myb protein expression: specific phosphorylation during the S phase of the cell cycle. Oncogene 12, 1855–1864

56. Delaglio, F., Grzesiek, S., Vuister, G. W., Zhu, G., Pfeifer, J., and Bax, A. (1995) NMRPipe: a multidimensional spectral processing system based on UNIX pipes. J Biomol NMR 6, 277–293

57. Lee, W., Tonelli, M., and Markley, J. L. (2015) NMRFAM-SPARKY: enhanced software for biomolecular NMR spectroscopy. Bioinformatics 31, 1325–1327

